# The Effect of THz Electromagnetic Field on the Conductance of Potassium and Sodium Channels

**DOI:** 10.1101/2024.08.27.609902

**Authors:** Zigang Song, Lingfeng Xue, Qi Ouyang, Chen Song

**Affiliations:** School of Life Sciences, Peking University, Beijing 100871, China; Center for Quantitative Biology, Academy for Advanced Interdisciplinary Studies, Peking University, Beijing 100871, China; School of Physics, Peking University, Beijing 100871, China; Peking-Tsinghua Center for Life Sciences, Academy for Advanced Interdisciplinary Studies, Peking University, Beijing 100871, China

**Keywords:** Ion channel, Potassium channel, Sodium Channel, Terahertz, Conductance

## Abstract

Ion channels are essential to various physiological processes and their defects are associated with many diseases. Previous research has revealed that Terahertz electromagnetic field can alter the channel conductance by affecting the motion of chemical groups of ion channels, and hence regulate the electric signals of neurons. In this study, we conducted molecular dynamics simulations to systematically investigate the effects of terahertz electromagnetic fields on the conductance of voltage-gated potassium and sodium channels, particularly focusing on the bound ions in the selectivity filters that have not been studied previously. Our results identified multiple new characteristic frequencies and showed that 1.4, 2.2, or 2.9 THz field increases the conductance of K_v_1.2, and 2.5 or 48.6 THz field increases the conductance of Na_v_1.5. The conductance-enhancing effects are specific to the frequencies and directions of the electric field, which are determined by the intrinsic oscillation motions of the permeating ions in the selectivity filter or certain chemical groups of the ion channels. The amplitude of the THz field positively correlates with the change in ion conductance. Therefore, this study demonstrates that THz fields can specifically regulate ion channel conductances, which may carry great potential in biomedical applications.

## 1 Introduction

Ion channels are integral membrane proteins that control the flow of ions across cellular membranes. They play crucial roles in numerous physiological processes, such as the transmission of neural signals and the initiation of muscle contractions, among other functions. These channels can be categorized based on their ion selectivity, distinguishing them as potassium, sodium, calcium, or chloride channels, or by their gating mechanisms, which include voltage-gated, ligand-gated, and mechanically-gated types. ^1–5^ Dysfunction of ion channels can lead to a variety of diseases, often referred to as channelopathies, including neurological disorders, cardiac arrhythmias, and muscular dystrophies. ^1^

Recently, a series of pioneering studies by Chang et al. demonstrated that terahertz (THz, 10^12^ Hz) electromagnetic fields can modulate the conductance of various ion channels, thereby influencing neuronal signaling. These findings are highly significant and may pave the way for new therapeutic approaches to treating channelopathies. For instance, an experimental study revealed that a 5.6 *μ*m wavelength (53.53 THz) field increased potassium current and modified action potential waveforms. ^6^ Another study showed that 8.6 *μ*m wavelength (34.88 THz) field enhanced auditory perception in guinea pigs. ^7^ Simulation studies have also indicated that a 51.87 THz field can increase the conductance of the potassium channel KcsA. ^8,9^ 42.55 THz field was found to alter the free energy profile of Ca^2+^ in the calcium channel CavAb. ^10^ 36.75 THz field was found to reduce the conductance of K_v_1.2 by decreasing channel stability, while 37.06 THz field had the opposite effect. ^11^ 15 THz field was reported to increase the conductance by enhancing the proportion of direct knock-on permeation. ^12,13^ A lower frequency of 0.1 THz field was also found to alter the energy profile and accelerate the permeation of K^+^ in KcsA. ^14^ A few other frequencies, such as 1 THz, were also examined. ^15^ These findings suggest that THz fields may have direct regulation effects on ion channels. In addition, terahertz waves have been explored for their ability to alter gene expression, detect tumors, and potentially reduce the side effects of drugs, ^16–19^ further demonstrating the therapeutic potential of the THz field.

Although the effects of THz fields on ion channels have been gradually recognized, most of the previous studies focused on the effects of THz fields on certain chemical groups of ion channels. Considering that there are usually well-defined ion binding sites within the selectivity filter of ion channels, one would expect that these bound ions should have well-defined oscillation motions, and the impact of THz fields on such motions remains largely uninvestigated. Therefore, this study aims to identify previously unknown frequencies that influence ion channel conductance, with a particular focus on those that directly affect the bound ions at the selectivity filter of ion channels.

We examined the structure and function of two well-studied ion channels: the voltage-gated potassium channel (K_v_) and the voltage-gated sodium channel (Na_v_). The K_v_ channel is a tetramer with each subunit featuring an N-terminal domain, six transmembrane helices (S1 ∼ S6), and a C-terminal domain. The voltage-sensing domain (VSD) is formed by S1 ∼ S4, while S5, S6, and the intervening pore loop constitute the pore region. The selectivity filter (SF) within the pore loop, characterized by the sequence TVGYGD, provides binding sites for K^+^ ions, named S0 ∼ S4. ^1,3–5,20–23^ The structure of a chimeric protein, K_v_1.2/2.1 (PDB: 2R9R), resolved in 2007, has been extensively studied and serves as a basis for our open-state K_v_1.2 model ^24–27^ (Figure 1A). The eukaryotic Na_v_ channel comprises an *α* subunit with four similar domains (DI ∼ DIV), each containing six transmembrane helices (S1∼ S6), along with several auxiliary *β* subunits. The SF is located at the pore loop between S5 and S6 and is formed by DEKA and EEDD residues. ^1,2,28–35^ Na_v_1.5 is a sodium channel expressed in cardiac muscle and encoded by *SCN5A*,^36,37^ whose partially-open structure was solved in 2021 (PDB: 7FBS).^35^ A recent study by Choudhury *et al*. proposed that the channel fully opens when the S6 segments in domains DI, DIII, and DIV or all four domains form partial *π*-helices. ^38^ Therefore, the Na_v_1.5 structure model with four manually-introduced *π* helices was used for this study (Figure 1B).

In this study, we performed molecular dynamics (MD) simulations to investigate the oscillation motions of bound ions in the SF of K_v_1.2 and Na_v_1.5, as well as the effects of THz frequencies on the K_v_1.2 and Na_v_1.5 channels. Our findings reveal that THz waves with specific frequencies, 1.4, 2.2, and 2.9 THz, can enhance the conductance of K_v_1.2, while 2.5 THz field increases that of Na_v_1.5, by directly affecting the motion of K^+^ or Na^+^ bound at the SF of the channels. Additionally, a 48.6 THz field increases Na_v_1.5 conductance by affecting the carboxyl groups (-COO^−^) at the SF. These results further suggest that THz fields could potentially be harnessed for therapeutic applications, offering a novel avenue for manipulating ion channel function.

**Figure 1.**
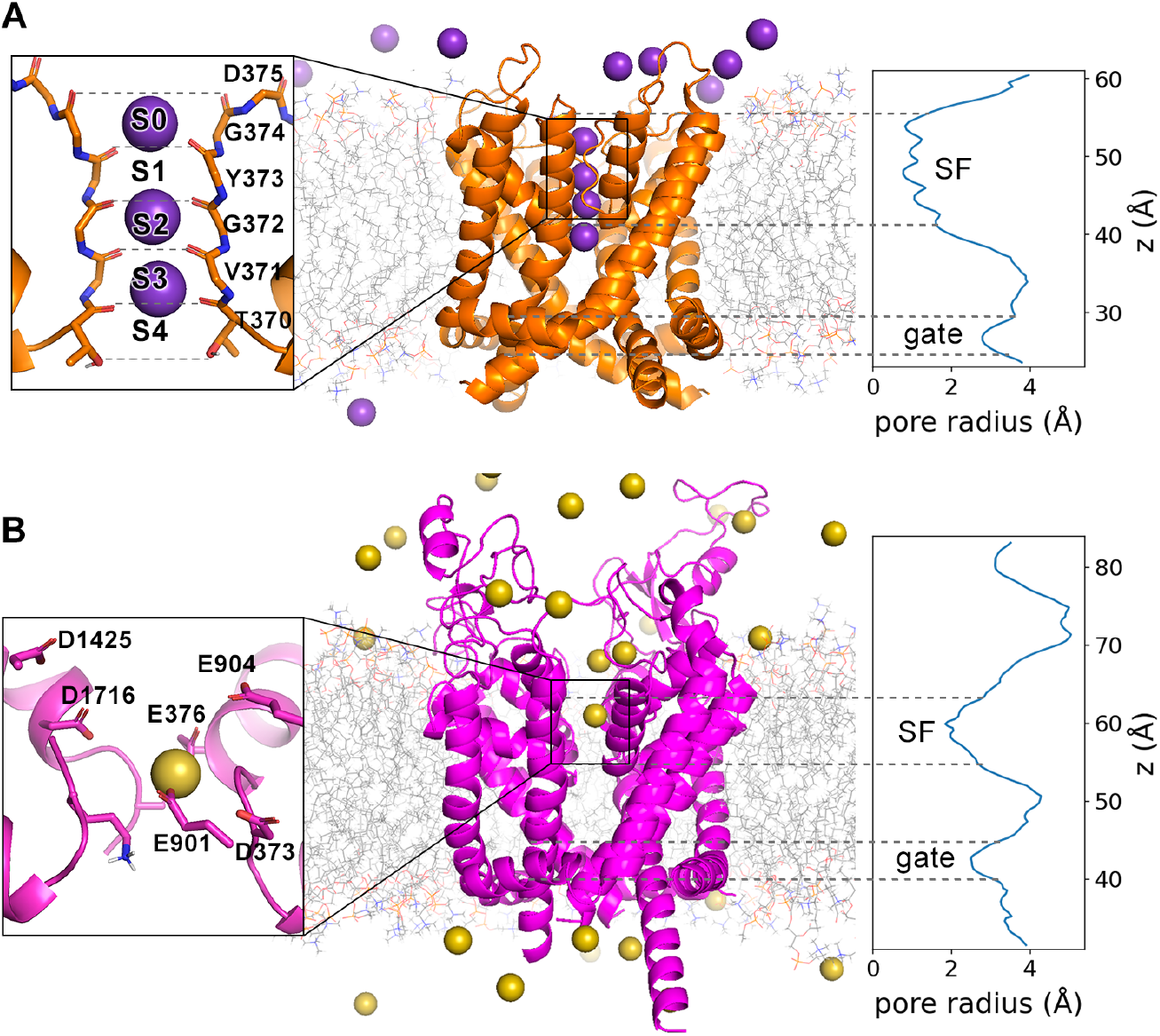
The simulation system. (A) K_v_1.2, (B) Na_v_1.5. Purple and yellow spheres represent K^+^ and Na^+^, respectively. Each left panel shows the structure of the selectivity filter (SF) with the bound ions, and each right panel shows the pore radius along the z-axis.

## 2 Results

### 2.1 Frequency analysis

To identify potential frequencies influencing ion motion within the channels, we performed a frequency analysis using Fast Fourier Transform (FFT) on the trajectories of ions and the chemical groups that form the ion binding sites within the selectivity filter. This analysis encompassed a frequency range from 0.1 to 100 THz.

Our findings identified distinct characteristic frequencies for both ions and chemical groups within the channels. For the potassium channel K_v_1.2, three potassium ions within the selectivity filter (SF), designated as K1 to K3, exhibited vibrational motions predominantly within the 1 to 6 THz range. Specifically, the peak frequencies for these ions in the z direction were 2.2, 2.9, and 1.4 THz, respectively, while their peak frequencies in the x or y directions were approximately 5.0 THz (Figure 2A).

**Figure 2.**
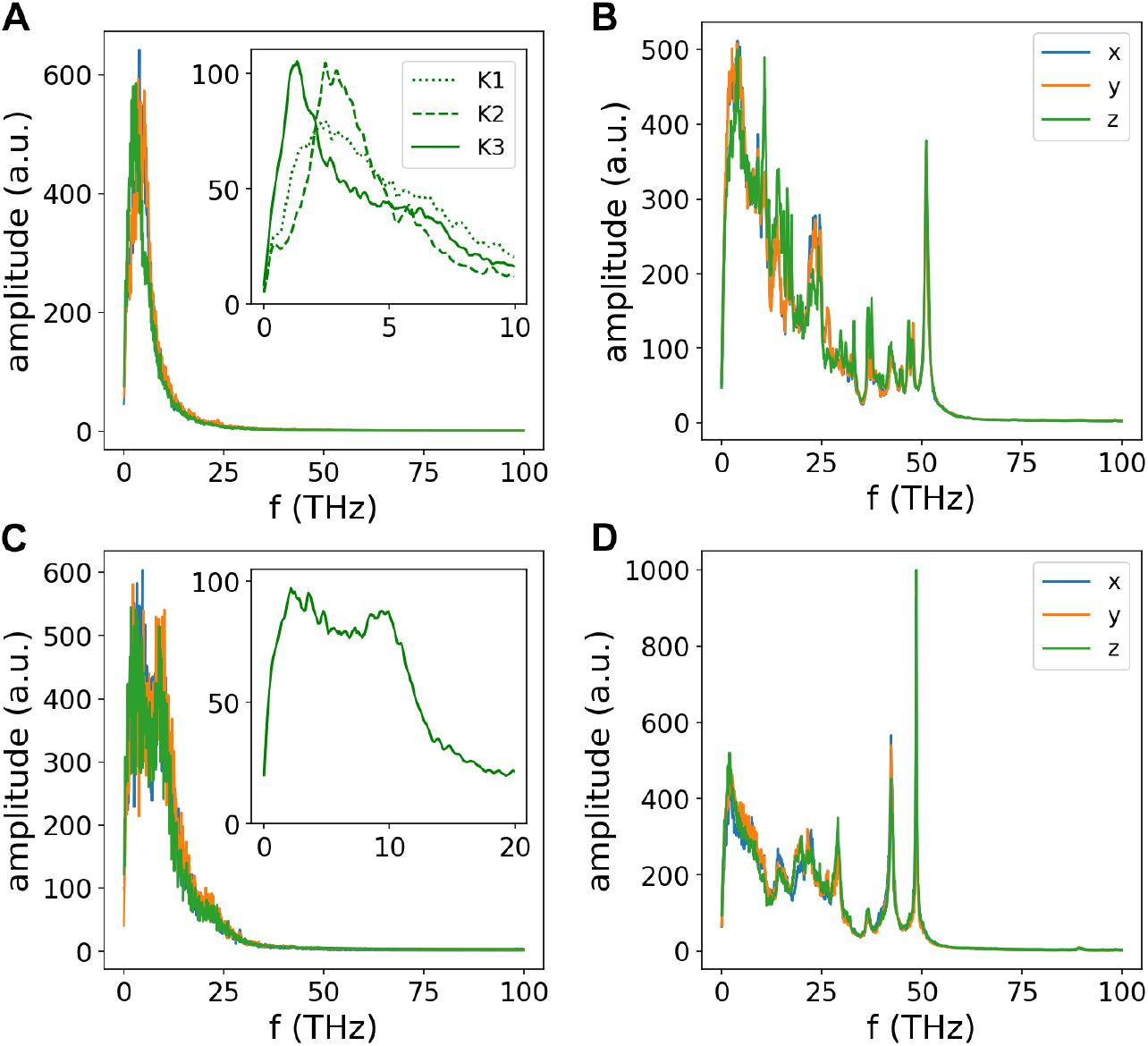
Frequency spectrum of (A) K^+^ ions within K_v_1.2, with the inset showing the z-direction spectrum of the three individual ions. (B) Oxygen atoms at the selectivity filter (SF) of K_v_1.2. (C) Na^+^ ion within Na_v_1.5, with the inset displaying its z-direction spectrum. (D) Carboxyl (-COO) groups at the SF of Na_v_1.5.

Additionally, we analyzed the carbonyl oxygen atoms of residues 370 to 375 in K_v_1.2, which form the ion binding sites. These atoms showed a peak frequency of 51.0 THz, corresponding to the stretching vibration mode of the C=O bond, which has been examined in the previous studies. ^8,9^ We also observed a peak at 10.8 THz, revealing a wiggling motion of the carbonyl oxygen atoms in the z-direction (Figure 2B).

For the sodium channel Na_v_1.5, the Na^+^ ion within the selectivity filter (SF) exhibited a broad frequency spectrum ranging from 1 to 12 THz. A prominent peak at 2.5 THz was associated with the ion’s motion in the x-z plane within the SF, while a smaller peak at 10.0 THz corresponded to motion in the y-z plane (Figure 2C). Additionally, we analyzed the carboxyl groups of glutamate and aspartate residues that interact with Na^+^ ions at the SF of Na_v_1.5 (residues 373, 376, 901, 904, 1425, and 1716). These groups exhibited peak frequencies at 48.6 THz and 42.5 THz, both linked to the stretching mode of the C-O bond. The 48.6 THz mode predominantly occurred in the x-z plane, while the 42.5 THz mode was observed in the x-y plane. Other minor peaks at lower frequencies were noted but are not discussed in this study (Figure 2D).

### 2.2 The effect of THz electromagnetic field

We initially performed control experiments without applying THz fields to establish a baseline for ion channel conductance. Conductance was measured by counting the number of ions passing through the channels over a specified time under certain transmembrane potentials.

For K_v_1.2, the simulated conductance was 12.6 ± 3.2 picosiemens (pS), consistent with the experimental range of 14 to 18 pS. ^20^ Throughout these simulations, ions permeated stably through the channel, with an average of three ions occupying the SF.

The simulated conductance of Na_v_1.5 was 39.1 ± 12.3 pS, exceeding but close to the experimental range of 10 to 20 pS. ^35,39^ This discrepancy may be due to the use of a more open structure in our simulations. Despite the elevated conductance, ion permeation through Na_v_1.5 remained stable, with an average of one sodium ion occupying the SF during the simulation period.

Subsequently, we applied electromagnetic fields at the observed frequencies to each ion channel system, focusing on the z-direction frequencies, as these align with the direction of ion permeation and are more likely to influence channel conductance. ^9^ The THz field was applied in the z direction with an amplitude of 0.8 V/nm, and channel conductance was measured under each condition. Additionally, we analyzed the ion distribution to determine whether the observed effects were associated with changes in ion distribution.

#### The effect of THz field on the potassium channel

For K_v_1.2, applying electromagnetic fields at frequencies of 1.4, 2.2, and 2.9 THz significantly increased channel conductance to 31.4 ± 6.6, 27.2 ± 6.5, and 30.8 ± 9.3 pS, respectively (n=6 per frequency, p<0.01). The effects were similar across these frequencies probably due to their proximity, with the 1.4 THz field producing the highest conductance, representing a 2.5-fold increase. In contrast, a 10.8 THz field decreased the conductance to 3.0 ± 1.0 pS (n=3, p<0.01), approximately one-quarter of the original value (Figure 3A,C).

**Figure 3.**
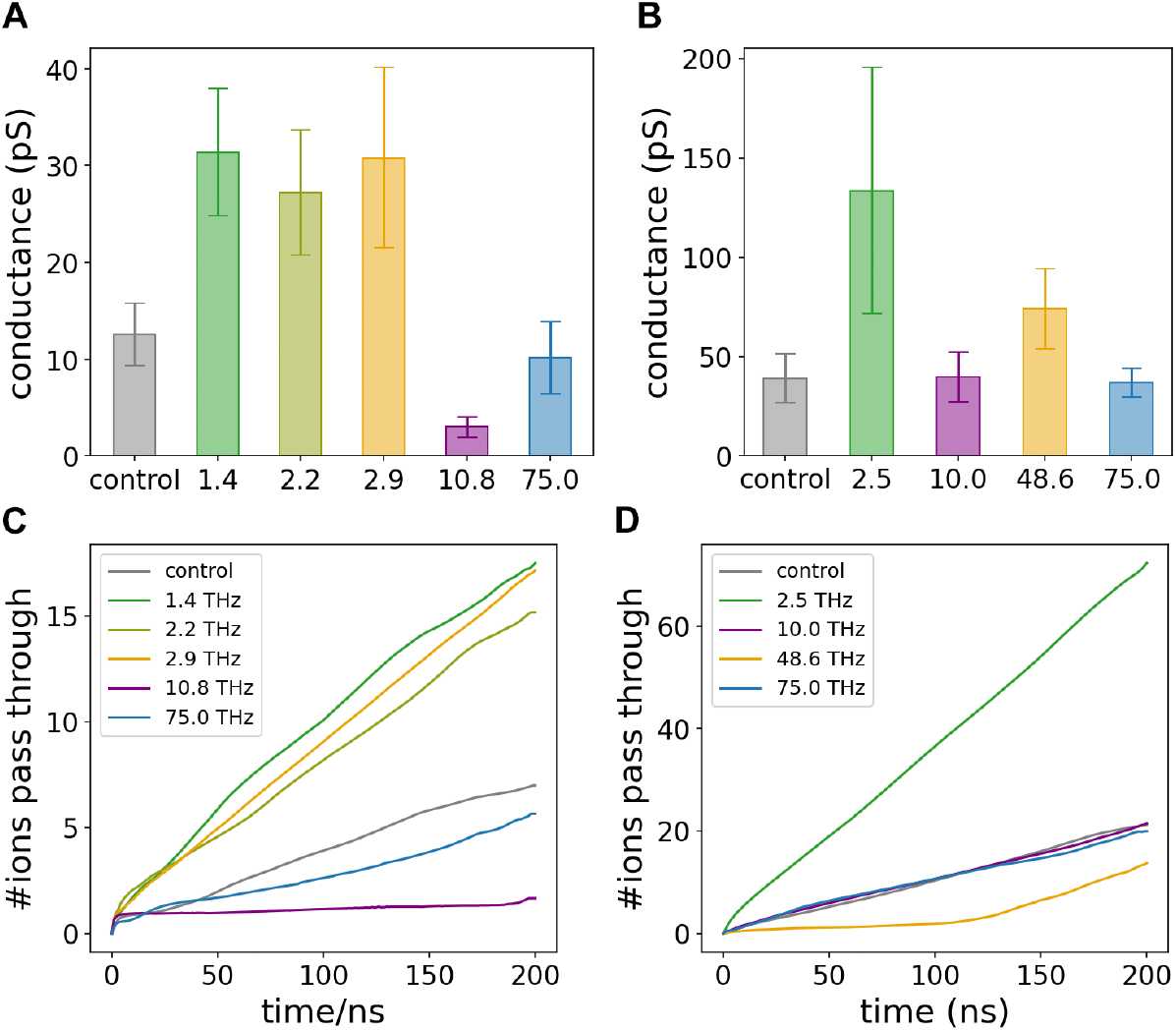
(A, B) The effect of THz electromagnetic fields at different frequencies on the conductance of K_v_1.2 (A) and Na_v_1.5 (B), respectively. The error bars represent the standard deviation across multiple simulation replicates. (C, D) The time-resolved average number of ions permeating through K_v_1.2 (C) and Na_v_1.5 (D), with data smoothed for clarity.

No changes in ion distribution were observed with the 1.4, 2.2, or 2.9 THz fields. However, the application of a 10.8 THz field altered the ion distribution. Typically, three K^+^ ions occupy the selectivity filter (SF) at S0, S2, and S3/S4 in the conducting state without the THz field. With the 10.8 THz field applied, the occupation state shifted to S1, S2/S3, and S4 (Figure 4A). This change in ion distribution likely led to a less conductive state and contributed to the observed decrease in conductance.

**Figure 4.**
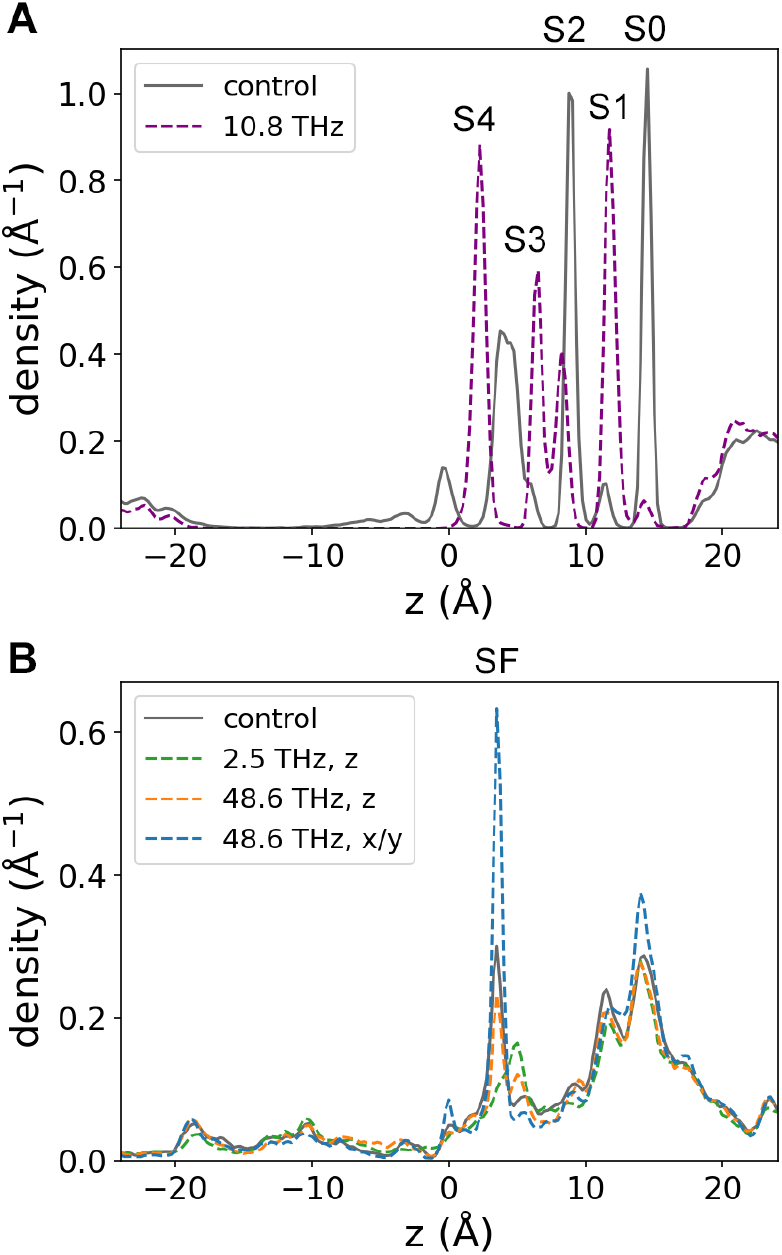
Ion distribution along the z-axis inside the channel, with or without the application of a THz field. (A) Distribution of K^+^ ions within K_v_1.2. (B) Distribution of Na^+^ ions within Na_v_1.5.

#### The effect of THz field on the sodium channel

For Na_v_1.5, the application of a 2.5 THz electromagnetic field dramatically reduced the residence time of Na^+^ ions and increased channel conductance to 133.5 ± 61.8 pS (n=6, p<0.05), representing a 3.4-fold increase compared to control conditions. The large fluctuation in conductance was due to an unusually low ion permeation count in one simulation trajectory, where only 10 ions permeated compared to the typical range of 70 to 110 ions in other runs. This anomaly occurred because a single Na^+^ ion occupied an atypical binding site, obstructing the passage of other ions. In contrast, applying a 10.0 THz field resulted in unchanged conductance at 39.7 ± 12.6 pS, indicating that not all types of enhanced motion induced by the THz field resulted in increased conductance (Figure 3B,D).

The 48.6 THz field also enhanced conductance, but this effect was preceded by a 140 ns delay during which conductance was initially low (Figure 3D). After extending the simulation to 600 ns, the conductance increased to 74.0 ± 20.2 pS (n=4, p<0.05), and further rose to 93.0 ± 22.6 pS when considering only the period beyond the initial delay (Figure S1). Given that the 140 ns delay is much shorter than most physiological processes, this effect is likely transient.

For both the 2.5 and 48.6 THz fields, a slight decrease in ion density at the SF was observed, which may be related to the increased permeation rate and reduced residence time of ions at the binding sites. (Figure 4B).

### 2.3 Simulations under non-specific conditions

#### The effect of non-specific frequencies

To assess whether the observed effects were specific to the peak frequencies, we performed additional simulations using non-specific frequencies. For this, we chose a 75.0 THz field, which is distant from the peak frequencies identified in our spectrum and thus less likely to cause interference.

No significant changes in channel conductance were observed when a 75.0 THz field was applied to each channel. The conductance values for K_v_1.2 and Na_v_1.5 were 10.2 ± 3.7 pS and 36.9 ± 7.4 pS, respectively (Figure 3). These values were consistent with the control measurements, suggesting that the changes in conductance observed in the above sections were specific to those selected frequencies and not due to a general effect of the THz field. Therefore, this control experiment reinforces the idea that the peak frequencies identified by FFT analyses have a specific effect on ion channels, rather than simply causing a general perturbation.

#### The effect of THz field in different directions

We further investigated the specificity of the THz field’s polarization direction on its effects by conducting simulations with the THz field applied in the x and y directions, in addition to the z direction.

For K_v_1.2, a 1.4 THz field significantly affected conductance when polarized in the z direction but had negligible effects when applied in the x or y directions (Figure 5A). Interestingly, although potassium ions showed a peak frequency of 5.0 THz in the x/y plane, applying an electric field at this frequency in the x or y directions did not significantly impact ion conductance. In contrast, applying the 5.0 THz field in the z direction increased ion conductance to 25.7 ± 2.1 pS (n=3, p<0.01), probably due to its proximity to the peak frequencies in the z direction (Figure 5A). This increase was less pronounced than that observed with the peak frequencies, further supporting the frequency specificity of THz fields.

Similarly, in the case of Na_v_1.5, applying a 2.5 THz field along the x or y direction did not result in significant changes in conductance (Figure 5B). Interestingly, when a 48.6 THz field was applied in the x or y direction, the conductance notably decreased to 10.1 ± 3.1 pS and 17.5 ± 2.3 pS, respectively (n=6 each, p<0.05) (Figure 5B). This decrease was accompanied by an increase in ion density at the SF (Figure 4B), indicating a tighter binding of Na^+^ ions to the SF of the channel, which likely caused the reduced conductance. These observations indicate that the influence of the THz field on ion channels is indeed specific to its polarization direction.

**Figure 5.**
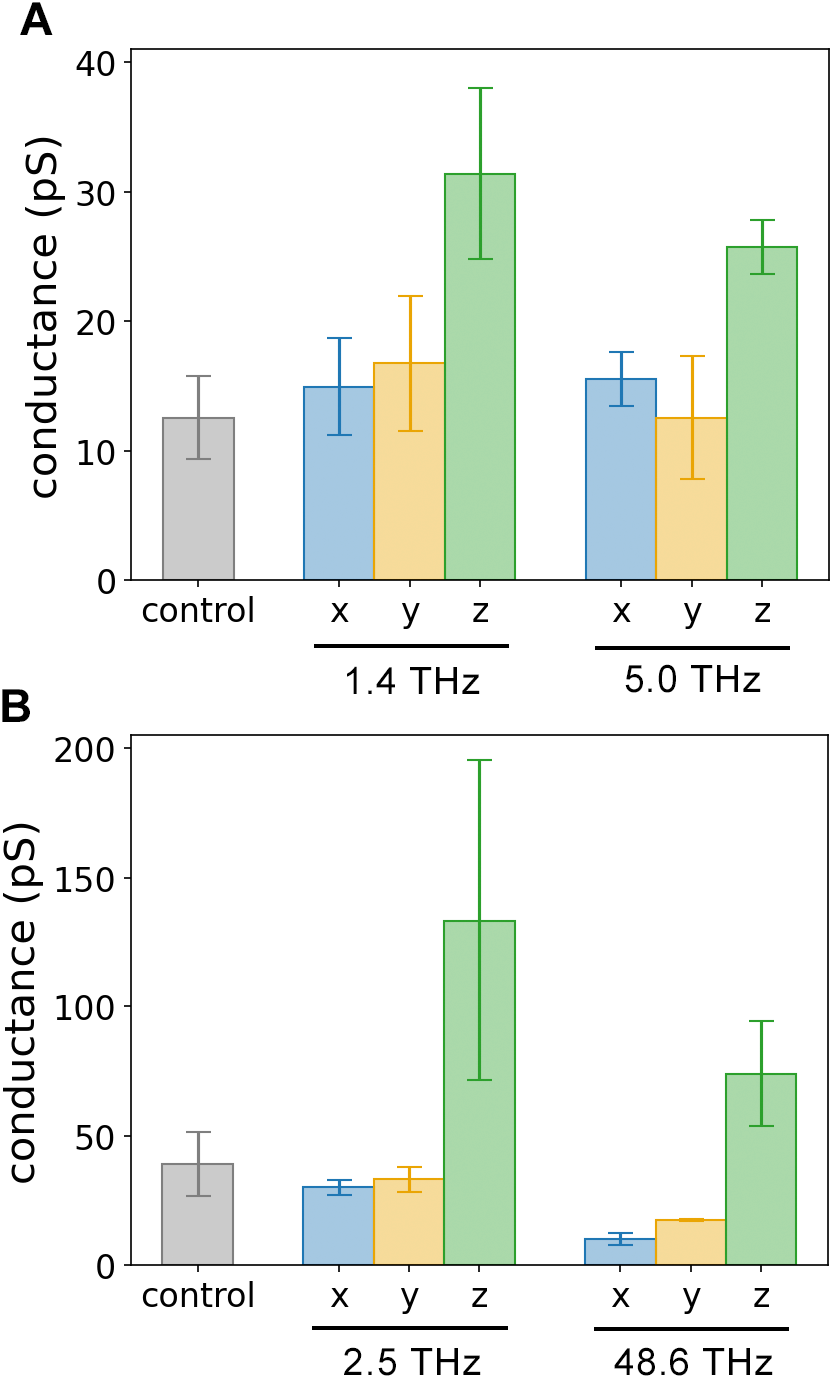
The effect of THz field in different directions on the conductances of (A) K_v_1.2 and (B) Na_v_1.5.

### 2.4 The effect of THz field with different amplitudes

We further investigated the effect of THz field amplitude by conducting simulations with varying amplitudes. For K_v_1.2, we applied a 1.4 THz electromagnetic field, while for Na_v_1.5, a 2.5 THz field was used. The amplitudes tested were 0.2, 0.4, and 0.8 V/nm. Both K_v_1.2 and Na_v_1.5 showed a positive correlation between the field amplitude and ion conductance (Figure 6).

The conductance of K_v_1.2 increased at all three amplitudes, with a linear relationship between amplitude and conductance, evidenced by a correlation coefficient of 0.84 (Figure 6A). In contrast, Na_v_1.5 showed a slightly different response. Conductance remained nearly unchanged at the lowest amplitude of 0.2 V/nm. A minor increase was noted at 0.4 V/nm, but a significant enhancement in conductance was observed only at the highest amplitude of 0.8 V/nm. This indicates a non-linear relationship between the THz field amplitude and the ion conductance, suggesting that a higher amplitude is required to significantly influence the conductance of Na_v_1.5 (Figure 6B).

**Figure 6.**
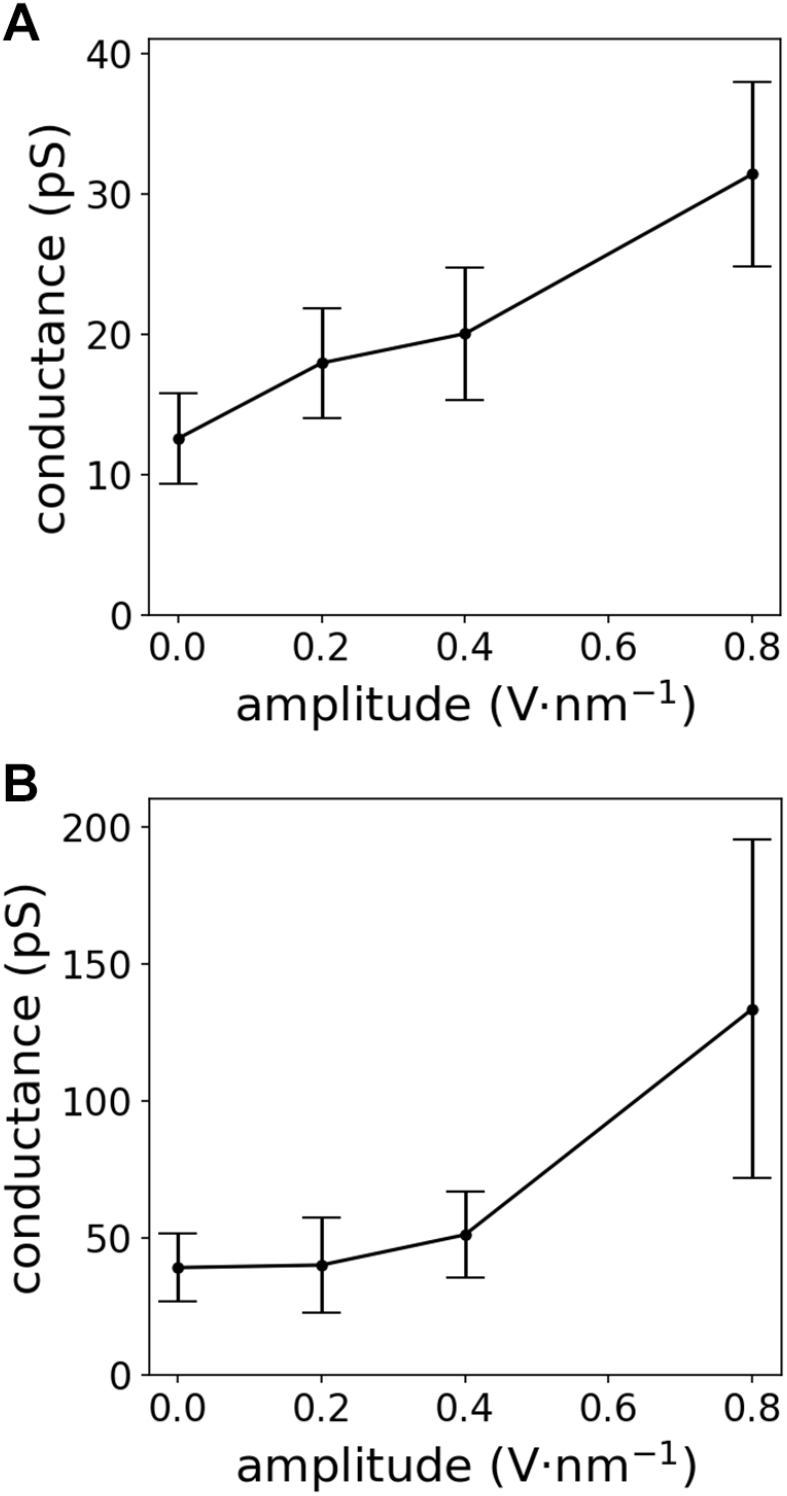
Relationship between THz field amplitude and conductance for (A) K_v_1.2 and (B) Na_v_1.5.

Consistently, we observed a negative correlation between the amplitude of the THz field and the residence time of ions at the SF, further confirming that the THz field influences ion motion and accelerates permeation (Figure S2). The residence time of K^+^ in K_v_1.2 decreased linearly with increasing amplitude, reaching a point where it was halved at an amplitude of 0.8 V/nm (Figure S2A). For Na_v_1.5, a similar trend was observed for the residence time of Na^+^ ions. The residence time started to decrease when the amplitude reached 0.4 V/nm, and reduced to one-fifth when the amplitude increased to 0.8 V/nm (Figure S2B).

## 3 Discussion

Despite numerous experimental and computational studies on the effects of THz fields on ion channels, it remained unclear at the molecular level whether the observed effects were specific or simply due to thermal influence. In our simulations, the system temperature was consistently maintained at 310 K with a high thermal coupling rate. No significant temperature changes were observed, with fluctuations of only 1.75 K in the K_v_1.2 system and 1.35 K in the Na_v_1.5 system. Moreover, if the effects of the THz field were purely thermal, they would not exhibit directional specificity. However, our simulations demonstrated that the THz field effects were both frequency- and direction-specific. Therefore, the observed changes in conductance in our simulations can be attributed to the non-thermal effects of the THz electromagnetic fields.

Our study demonstrates that the influence of THz electromagnetic fields on ion channels is highly specific to both frequency and polarization direction. THz fields resonating at the frequencies of potassium ions within K_v_1.2 (1.4, 2.2, and 2.9 THz) and sodium ions within Na_v_1.5 (2.5 THz) directly enhance channel conductance by accelerating ion motion along the permeation direction (z-axis). In contrast, a 10.8 THz field, resonating with the carbonyl oxygen atoms of the selectivity filter, alters ion distribution within K_v_1.2, resulting in decreased conductance. These findings suggest that ion channel conductance can be modulated through different mechanisms, underscoring a multifaceted approach to controlling molecular behavior. While previous research focused on frequencies associated with the protein motions, which influence ion motion indirectly, our study shifts the focus to frequencies that resonate directly with the permeating ions at the SF, yielding more pronounced effects.

Our study demonstrates that ion channel function can be modulated by specific THz fields, opening new avenues for applications in biomedical fields, particularly in the treatment of channelopathies. The non-invasive and non-destructive nature of THz waves, combined with their ability to be precisely directed and focused, presents a promising therapeutic modality. Moreover, this approach could be extended to identify specific frequencies that affect other biomolecules, offering a versatile tool for molecular manipulation.

Despite these promising findings, our study has some limitations. The simulated electric field amplitude of 0.8 V/nm is significantly higher than what might be safely applied in practical scenarios, and the simulation duration may not fully capture the effects over longer periods. Therefore, further experimental studies are essential to validate the impact of THz fields on ion channels and their effects on cellular behavior at a macroscopic level. These future investigations should focus on refining the intensity and duration of exposure to ensure both safety and efficacy in real-world applications.

## 4 Methods

### 4.1 Model systems

The K_v_1.2 model was built with the open-state structure of a chimeric protein K_v_1.2/2.1 (PDB: 2R9R). Constructed from rat K_v_1.2 and K_v_2.1, this protein has an identical pore region as K_v_1.2, and produces a similar current to K_v_1.2. ^24^ The pore region (residues 309 ∼ 417 of each subunit) and the K^+^ within were selected and embedded in 1-palmitoyl-2-oleoyl-glycero-3-phosphocholine (POPC) bilayer to mimic the cellular membrane environment. The system was solvated with an aqueous solution containing 150 mM KCl. The channel’s orientation was set so that its axis aligned along the z direction, with the membrane parallel to the x-y plane. The TIP3P model was used for water molecules. The simulation system was prepared using CHARMM-GUI. ^40^

The Na_v_1.5 model was built using the open-state structure with 4 *π*-helices, constructed by Choudhury *et al*. ^38^ The pore region (residues 236 ∼ 430, 825 ∼ 945, 1320 ∼ 1478, 1643 ∼ 1777) was selected and embedded in a POPC bilayer. The system was solvated in a solution containing 150 mM NaCl.

To confirm that the channels were in the open state, we calculated the pore radius of each channel using a PyMol plugin CAVER 3.0.3. ^41^

### 4.2 MD simulations

We performed molecular dynamics (MD) simulations using GROMACS 2022.3, ^42^ augmented with an in-house developed plugin that enabled the addition of a constant electric field in each spatial direction (x, y, or z). This capability allowed us to simulate ion permeation in the presence of both an oscillating electric field and a uniform electric field along the channel axis.

The simulations were conducted under various conditions (Table 1), each with 3 ∼ 6 replicates. The control and specific conditions were simulated with six replicates, while non-specific conditions were repeated three times. A total of 24 *μ*s simulation was performed in this study.

**Table 1.** Simulation systems, conditions, and replicates.

| System | Frequency [THz] | Direction | Amplitude [$\text{V nm}^{-1}$] | Time [ns] | Replicates |
| --- | --- | --- | --- | --- | --- |
| $\text{K}_v1.2$ | - | - | - | 200 | 6 |
|  | 1.4 | x | 0.8 | 200 | 3 |
|  | 1.4 | y | 0.8 | 200 | 3 |
|  | 1.4 | z | 0.2 | 200 | 6 |
|  | 1.4 | z | 0.4 | 200 | 6 |
|  | 1.4 | z | 0.8 | 200 | 6 |
|  | 2.2 | z | 0.8 | 200 | 6 |
|  | 2.9 | z | 0.8 | 200 | 6 |
|  | 5.0 | x | 0.8 | 200 | 3 |
|  | 5.0 | y | 0.8 | 200 | 3 |
|  | 5.0 | z | 0.8 | 200 | 3 |
|  | 10.8 | z | 0.8 | 200 | 3 |
|  | 75.0 | z | 0.8 | 200 | 3 |
| $\text{Na}_v1.5$ | - | - | - | 200 | 6 |
|  | 2.5 | x | 0.8 | 200 | 3 |
|  | 2.5 | y | 0.8 | 200 | 3 |
|  | 2.5 | z | 0.2 | 200 | 6 |
|  | 2.5 | z | 0.4 | 200 | 6 |
|  | 2.5 | z | 0.8 | 200 | 6 |
|  | 10.0 | z | 0.8 | 200 | 6 |
|  | 48.6 | x | 0.8 | 200 | 6 |
|  | 48.6 | y | 0.8 | 200 | 6 |
|  | 48.6 | z | 0.8 | 600 | 4 |
|  | 75.0 | z | 0.8 | 200 | 3 |

The CHARMM36m force field ^43,44^ and periodic boundary condition (PBC) were employed for the simulation. Leap-frog algorithm was used with a 2 fs time step. Temperature was maintained at 310 K using the Nose-Hoover thermostat, ^45,46^ and pressure was kept constant at 1 atm using the Crescale barostat. ^47^ Particle-Mesh Ewald (PME) was used for long-range electrostatic calculations. ^48^ Van der Waals forces were considered up to a cutoff of 1.2 nm, with a switching function applied from 1.0 nm to smooth the transition.

To maintain the open state of the channels, a position restraint of 100 kJ/(mol·nm^2^) was applied to the protein backbone, except for the selectivity filter region (residues 370 ∼ 375 for K_v_1.2, and residues 372 ∼ 376, 900 ∼ 904, 1420 ∼ 1425, 1712 ∼ 1716 for Na_v_1.5). A constant electric field *E*_0_ was applied along z direction to establish a +450 or -450 mV transmembrane potential, calculated as the product of *E*_0_ and the length of the box along z direction *l*_*z*_ ^26^

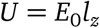

The ratio of electric field to magnetic field in electromagnetic waves equals the light speed *c*. Given that the velocities of charged particles in the system are far lower than *c*, the impact of the magnetic component is negligible. ^10^ Consequently, only the electric component was considered in our simulations. The total electric field applied to the system was therefore

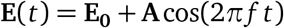

Where *A* and *f* denote the amplitude and frequency of the THz field.

### 4.3 Frequency analysis

After the equilibration of each system, a 1-ns simulation was performed for the frequency analysis. The same simulation setup was adopted as above, except that no restraints or electric field were applied. We analyzed the velocities of ions and relevant chemical groups and performed the Fast Fourier Transform (FFT) to obtain their frequency spectra.

### 4.4 Calculation of conductance

The conductance of a channel was calculated from the number of ions passing through in the simulation time under the transmembrane potential:

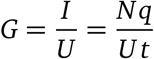

Where *N* is the number of ions passing through in time *t, I* and *U* denote the current and transmembrane potential, and *q* is the charge of each ion.

## 5 Acknowledgements

The MD simulations were performed on the computing platform of the Center for Life Sciences at Peking University and the National Supercomputer Center in Tianjin.

## 6 Data availability statement

The MD plugin can be found on GitHub (https://github.com/ComputBiophys/GromacsPlugin). The simulation data are available upon reasonable request.

