## Supplementary Information for "The Effect of THz Electromagnetic Field on the Conductance of Potassium and Sodium Channels"

### 1 Supplementary Figures

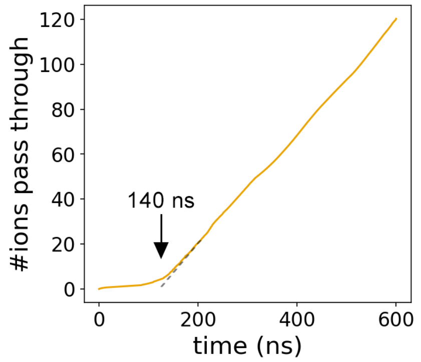


**Figure S1:** The average number of ions passing through $Na_v1.5$ when the 48.6 THz field was applied in the z-direction. The data was smoothed for clarity.

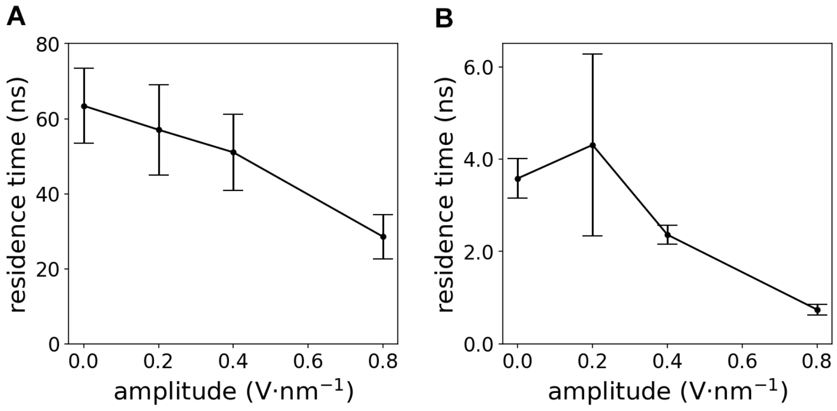


**Figure S2:** Relationship between THz field amplitude and ion residence time in the selectivity filter for (A) $K_v1.2$ and (B) $Na_v1.5$.
